# Phylogenomics of globally spread Clonal Groups 14 and 15 of *Klebsiella pneumoniae*

**DOI:** 10.1101/2022.08.30.505806

**Authors:** Carla Rodrigues, Val F. Lanza, Luísa Peixe, Teresa M. Coque, Ângela Novais, the Epidemiological Markers Study Group (ESGEM)

## Abstract

The increasing worldwide spread of multidrug-resistant (MDR) *Kp* is largely driven by high-risk sublineages, some of them well-characterised such as Clonal Group (CG) 258, CG147 or CG307. MDR *Kp* Sequence-Type (ST) 14 and ST15 have been described worldwide causing frequent outbreaks of CTX-M-15 and/or carbapenemase producers. However, their phylogeny, population structure and global dynamics remain unclear. Here, we clarify the phylogenetic structure and evolvability of CG14 and CG15 *Kp* by analysing the CG14 and CG15 genomes available in public databases (n=481, November 2019) and *de novo* sequences representing main sublineages circulating in Portugal (n=9). Deduplicated genomes (n=235) were used to infer temporal phylogenetic evolution and to compare their capsular locus (KL), resistome, virulome and plasmidome using high-resolution tools.

Phylogenetic analysis supported independent evolution of CG14 and CG15 within two distinct clades and 4 main subclades which are mainly defined according to the KL and the accessory genome. Within CG14, two large monophyletic subclades, KL16 (14%) and KL2 (86%), presumptively emerged around 1937 and 1942, respectively. Sixty-five percent of CG14 carried genes encoding ESBL, AmpC and/or carbapenemases and, remarkably, they were mainly observed in the KL2 subclade. The CG15 clade was segregated in two major subclades. One was represented by KL24 (42%) and KL112 (36%), the latter one diverging from KL24 around 1981, and the other comprised KL19 and other KL-types (16%). Of note, most CG15 genomes contained genes encoding ESBL, AmpC and/or carbapenemases (n=148, 87%) and displayed a characteristic set of mutations in regions encoding quinolone resistance (QRDR, GyrA83F/GyrA87A/ParC80I). Plasmidome analysis revealed 2463 plasmids grouped in 27 predominant plasmid groups (PG) with a high degree of recombination, including particularly pervasive F-type (n=10) and Col (n=10) plasmids. Whereas *bla*_CTX-M-15_ was linked to a high diversity of mosaic plasmids, other ARGs were confined to particular plasmids (e.g. *bla*_OXA-48_-IncL; *bla*_CMY/TEM-24_-IncC). This study firstly demonstrates an independent evolutionary trajectory for CG15 and CG14, and suggests how the acquisition of specific KL, QRDR mutations (CG15) and ARGs in highly recombinant plasmids could have shaped the expansion and diversification of particular subclades (CG14-KL2, CG15-KL24/KL112).

**IMPORTANCE:** *Klebsiella pneumoniae* (*Kp*) represents a major threat in the burden of antimicrobial resistance (AMR). Phylogenetic approaches to explain the phylogeny, emergence and evolution of certain multidrug resistant populations have mainly focused on core-genome approaches while variation in the accessory genome and the plasmidome have been long overlooked. In this study, we provide unique insights into the phylogenetic evolution and plasmidome of two intriguing and yet uncharacterized clonal groups (CGs), the CG14 and CG15, which have contributed to the global dissemination of contemporaneous β-lactamases. Our results point-out an independent evolution of these two CGs and highlight the existence of different clades structured by the capsular-type and the accessory genome. Moreover, the contribution of a turbulent flux of plasmids (especially multireplicon F type and Col) and adaptive traits (antibiotic resistance and metal tolerance genes) to the pangenome, reflect the exposure and adaptation of *Kp* under different selective pressures.

## INTRODUCTION

*Klebsiella pneumoniae* (Kp1, hereafter *Kp*) is one of the urgent threats for the emergence and spread of antimicrobial resistance (AMR) (1, 2), and one of the top six human pathogens leading to deaths attributable to AMR in 2019 (3). The increase of nosocomial infections caused by multidrug resistant (MDR) *Kp* and community-acquired infections caused by hypervirulent strains represents major a Public Health problem (4, 5). Recent advances in the population structure of *Kp* using core-genome multilocus sequence typing (cgMLST) or comparative analysis of whole genome sequences (WGS) revealed the predominance of a few clonal groups (CG) associated with antibiotic resistance (CG15, CG29, CG147, CG101, CG231, CG258, and CG307) (5, 6). However, to date, high-resolution genomic analysis is available only for a few of these predominant MDR (CG258, ST307, CG101, CG147) (7–10). *Kp* Sequence-Type (ST) 14 and ST15 strains represent 2.1% and 5.2% of the publicly available genomes (accessed in July 2020), respectively, and are frequently ESBL or carbapenemase producers that confer resistance to different antibiotics and are involved in hospital outbreaks worldwide (11, 12). ST15 and ST14 were initially considered to be highly related based on the single point mutation at the *infB* allele of the MLST scheme (http://bigsdb.pasteur.fr/klebsiella/klebsiella.html) (13). High resolution genomic analysis using small sample sizes yield inconsistent results. Whereas some studies referred to them as a single clonal group (CG15) (14, 15), others suggested CG15 and CG14 constitute two different CGs (6, 16, 17). Furthermore, recombination-free maximum-likelihood phylogeny revealed a deep branching structure with two major clades comprising several clusters of isolates with distinct capsular (KL) types (15). However, a clear correlation and phylogenetic analysis is still missing.

As other MDR *Kp* CGs, CG14 and CG15 are deep-branching lineages with a low nucleotide divergence (<0.5%) that largely vary on their accessory genome. Genomic variation is introduced by large homologous recombination events involving the capsule locus or via exchange of plasmids, phages or integrative conjugative elements (ICEs) (5, 15). The influence of the mobilome, and especially of the plasmidome, in shaping the evolutionary routes and adaptation of nosocomial pathogens has been scarcely explored (16, 18) mainly due to difficulties in puzzling short read sequencing data or performing long read sequencing in large collections of isolates (19–22). For this reason, most studies address plasmid diversity by replicon typing (23), which fail to establish whole plasmid entities. Moreover, a comprehensive analysis of the clonal diversification considering both core and accessory genome is lacking.

This study analyses the diversity and evolvability of CG14 and CG15 *Kp* genomes focusing on both its core and accessory genome (antimicrobial resistance, capsular polysaccharide locus and plasmidome) using high resolution tools (24, 25).

## RESULTS

### Phylogenomic analysis of global *Kp* CG15 and CG14

We performed a detailed phylogenomic analysis of 235 non-duplicated CG14 (n=65) and CG15 (n=170), including publicly available genomes (n=226) that met our quality criteria and genomes (n=9) *de novo* sequenced in this study (see material and methods for details) (**Table S1, Table S3**). Figure 1 shows the maximum-likelihood phylogenetic tree based on the concatenation of 4,420 core genes (representing ~80% of a *Kp* genome) which grouped the genomes according to the CG and the capsular type into two distinct clades (CG14 and CG15) and their subclades (KL2 and KL16 within CG14; and KL24, KL112, and a group of variable K-types including KL19 within CG15). The SNPs median distance between CG14 and CG15 genomes was 2452 SNPs/Mb (ranging between 7,500 and 12,143 SNPs), being much lower between the genomes of each clade, especially those of the CG15 (CG14: 541-2,328 SNPs/Mb; CG15: 252-1,084 SNPs/Mb) (**Table S2, Figure S1**).

**Figure 1.**
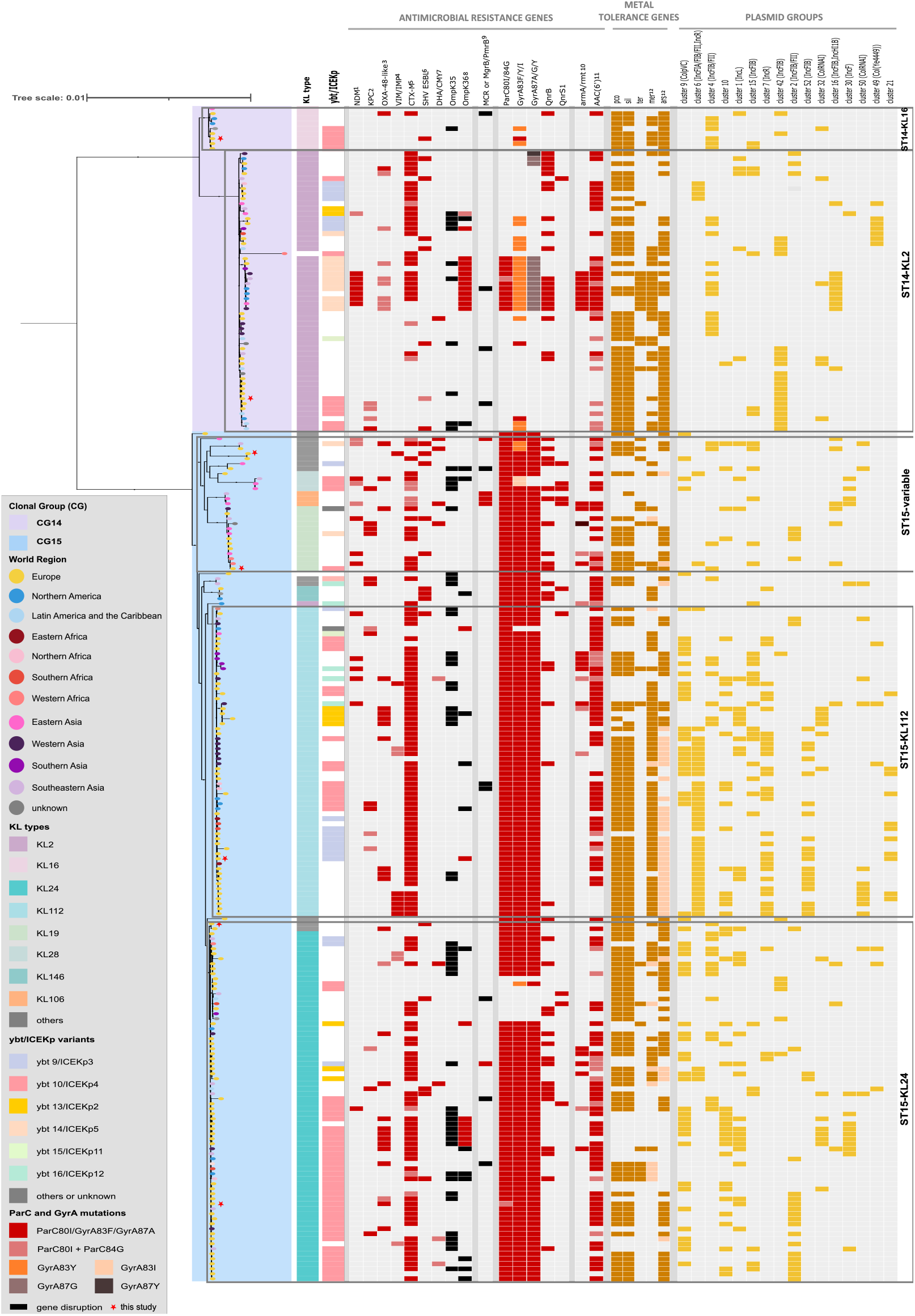
Phylogenetic structure of *K. pneumoniae* CG14 and CG15. Maximum-likelihood tree (model GTR+F+ASC+R4) inferred from 35,783 SNPs extracted from the alignment of 4,420 core genes, and rooted using two outgroups [*K. pneumoniae* ST540 Kpn0019 (accession number: SRR2098710) and a *K. pneumoniae* ST101 Kp_Goe_33208 (GCF_001902435.1)]. Branch lengths represent the number of nucleotide substitutions per site (scale, 0.01 substitution per site). The two main clades correspond to CG14 and CG15 and are shaded accordingly (see key). The branch tips are colored by the world region of isolation (see key). Capsular locus (KL) type, yersiniabactin-carrying ICEKp elements and GyrA and ParC mutations are colored according to their variants (see key). Antimicrobial-resistance determinants, metal tolerance clusters and plasmid groups are indicated by coloured rectangles when present. Gene disruption is represented by black rectangles. 1, dark pink indicates NDM-1 and light pink indicates other NDM-1 variants; 2, dark pink indicates KPC-2 and light pink indicates KPC-3; 3, dark pink indicates OXA-48 and light pink indicates other OXA-48-like variants; 4, dark pink indicates VIM whereas light pink stands for IMP; 5, dark pink indicates CTX-M-15 and light pink indicates other CTX-M variants; 6, dark pink indicates SHV-ESBL whereas light pink stands for TEM-ESBL; 7, dark pink indicates DHA-1 whereas light pink stands for CMY; 8, dark pink stands for the OmpK36GD mutation, light pink stands for other OmpK36 mutations; 9, dark pink indicates MCR presence whereas black indicates MgrB or PmrB disruption; 10, dark pink indicates ArmA and light pink indicates Rmt presence; 11, dark pink indicates the presence of the AAC(6’)-Ib-cr variant and light pink the presence of other AAC(6’) variants. 12, light yellow indicates that the cluster is not complete.

Besides the KL-type diversity, CG14 and CG15 clades also varied greatly in their accessory genome including antibiotic resistance genes (ARG), virulence gene determinants (e.g. yersiniabactin), metal tolerance genes (MTG) and plasmid replicons, as explained below (**Figure 1, Table S1**). Ninety-two percent of the genomes (n=217/235) carried O1 antigen type (92% variant O1v1, 8% O1v2) while the occurrence of other antigen types was sporadic (3 O2, 1 O4). All CG15 and CG14 genomes carried genes encoding type 3 fimbrial gene cluster (*mrk*), an iron ABC transporter (*kfu*), and a complete copy or remnants of the *kpi* chaperone-usher pili system (26), showing its association not only with CG15 but also with CG14. The putative virulence genes *rmpA2*, the aerobactin (*iuc*) and salmochelin (*iro*) siderophores were rarely observed. Contrary, the presence of yersiniabactin siderophore (*ybt*), which is usually mobilizable through an integrative conjugative element (ICE), was variable (51%). The lineage *ybt*10/ICEKp4 (57%) was the most frequent one and was randomly distributed among the subclades of the CG15 and CG14 (**Figure 1, Table S1**).

### Phylogenetic structure and diversity within the *Kp* CG14

The 65 CG14 genomes analysed were structured in two subclades representing KL2 (86%; 28-2346 SNPs/Mb) and KL16 (14%; 124-464 SNPs/Mb) (**Figure 1**). A phylogenetic inference (root-to-tip regression analysis: *R^2^* =0.1136; **Figure S2B, Figure S3**) suggested a relative time-scale emergence of the two CG14 subclades. Accordingly, the evolutionary rate within CG14 was estimated at 4.92×10^-7^ substitutions/site/year [95% highest posterior density (HPD), 3.70×10^-7^-6.16×10^-7^] corresponding to 2.1 SNPs/genome/year (95% HPD, 1.3-2.6). The ancestor of CG14 was estimated around 1836, although with a large uncertainty (95% HPD, 1762-1901), and the two predominant subclades, KL16 and KL2, emerged around 1937 (95% HPD, 1909-1963) and 1942 (95% HPD, 1926-1958), respectively.

CG14 *Kp* clinical isolates have been identified in many countries (45% in Europe, 27% in Asia, 21% in America) between 1980 and 2018 (**Figure S4**). The *Kp* ST14 strains from Portugal represented clones that contribute to the dissemination of genes encoding SHV-106 (ST14-KL16) or TEM-24 (ST14-KL2) in the early 2000’s (27, 28) (**Table S3**). One additional KL2 genome collected in Portugal in 1980 (Kp45) lacks canonical ARGs and represents the oldest isolate recovered from a human infection described to date. Genes encoding ESBL, AmpC and/or carbapenemases (mainly *bla*_CTX-M-15_, *bla*_OXA-232_, *bla*_TEM-24_, *bla*_CTX-M-14/-36_ and/or *bla*_NDM-1_) were observed in 65% of CG14 genomes and, remarkably, they were more frequently detected within the KL2 subclade (22% in KL16 *versus* 71% in KL2, *p*=0.0042). Additionally, variable mutations in genes coding for quinolone resistance determining regions (QRDR) were also observed in 42% (27/65) of the CG14 genomes (mainly GyrA-83Y, GyrA-87G, ParC-80I) (**Figure 1, Table S1**). SHV-28 was the most frequent narrow-spectrum SHV β-lactamase identified (57%, n=37/65).

### Phylogenetic structure and diversity within the *Kp* CG15

The 170 CG15 *Kp* genomes were segregated in differently populated subclades. The largest one included genomes carrying KL types 24 (42%; 72/170; 26-923 SNPs/Mb) and 112 (36%; 62/170; 24-31 SNPs/Mb). The other major subclade comprised genomes with KL19 (48%; 13/27; 31-404 SNPs/Mb) and other diverse KL-types (4 KL28, 2 KL62, 2 KL64, 3 KL106, 2 KL110, 1 KL131) (**Figure 1, Figure S1, Table S2**). The time-scale phylogenetic inference (root-to-tip regression analysis: *R^2^* = 0.0781, **Figure S2C, Figure S5**) suggests an evolutionary rate within CG15 estimated at 6.19×10^-7^ substitutions/site/year (95% HPD, 4.83×10^-7^-7.56×10^-7^) corresponding to 2.6 SNPs per genome per year. The ancestor of CG15 was estimated around 1858 although with a large uncertainty (95% HPD, 1763-1931).

The main subclades emerged both around 1960. One of them (95% HPD, 1950-1970) comprised one branch of KL24 isolates with wild-type *gyrA* and *parC* genes, and a second branch of KL24 isolates with GyrA83F, GyrA87A and ParC80I mutations that emerged around 1981 (1970-1990). The latter was subsequently split around 1987 (1980-1993) in one subclade dominated by the KL112 capsular locus and a small subclade of KL24 (n=5) suggesting recombination events and further selection of the KL112 capsular type (**Figure S5)**. The other subclade emerging around 1960 included one branch with a few genomes with KL106 and a second branch comprising genomes with KL19 locus that, apparently, emerged more recently (1988-2000). All these also carried the same QRDR mutations. Genomes carrying KL106, KL110 or KL39 were reported from China, Thailand and Portugal, respectively, and might correspond to local adaptive events (**Table S3**).

The CG15 genomes analysed in this work corresponded to isolates identified between 1980 and 2017, mostly collected in Europe (59%, 100/170), and from the EuScape study (https://pathogen.watch/collections/all?searchText=euscape) (65%, 65/100). Those collected in Portugal and sequenced *de novo* in this work represent predominant lineages in Europe, linked to CTX-M-15 (KL112) or CTX-M-15+OXA-48 production (27, 29, 30) (**Figure S4, Table S3**). Considering the 170 CG15 genomes analysed, most (n=148; 87%) carry genes encoding ESBL, AmpC and/or carbapenemases. CTX-M-15 is the predominant enzyme (68%; 116/170), which was significantly enriched in the KL24/KL112 large subclade (64% vs. 92%, *p*=0.0004). Carbapenemases were detected in 38% (n=64/170) of CG15, being OXA-48-like the most frequently detected carbapenemase (41%; 26/64). A higher diversity of genes encoding ESBLs (CTX-M, SHV-2/-12, DHA-1, VIM-34, NDM-19) were observed in genomes carrying KL19 and other diverse capsular loci. Similarly to CG14, *bla*_SHV-28_ was identified in most (91%; 155/170) CG15 isolates (**Table S1**).

### The accessory genome of *Kp* CG15 and CG14

Using the Pangenome Analysis Toolkit (PATO) to define and represent the accessory genome (see material and methods for details), we identified a common and specific proteoma for CG14 and CG15 and main subclades thereof (>80% of the genomes and with >80% identity, p-value < 0.001) (**Figure 2**). From the proteins enriched in each of the CG (>80% of the genomes, p-value < 0.001), 52% (n=109/211) of them were annotated as hypothetical proteins (**Table S4**).

**Figure 2.**
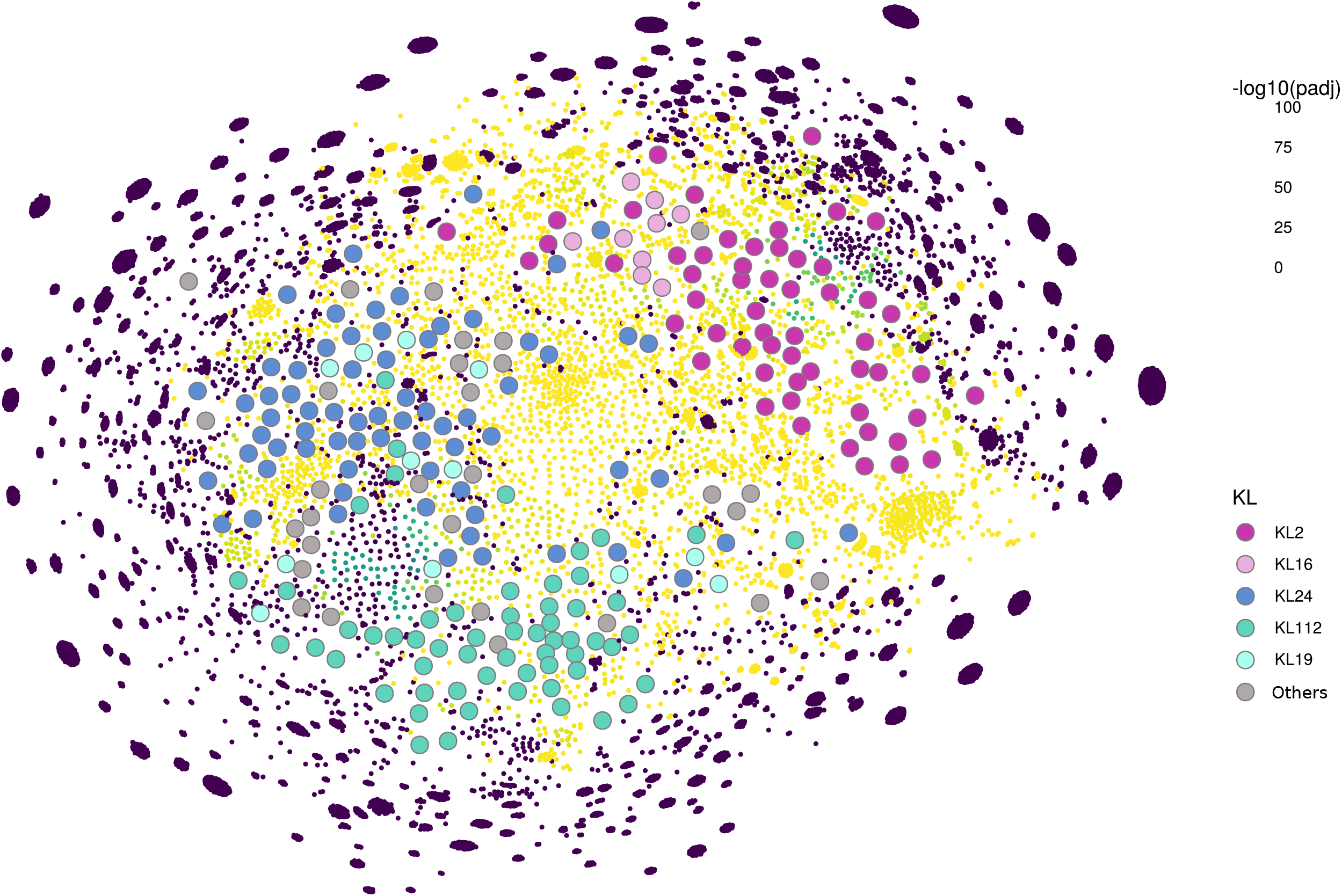
Accessory Genome Network (AcCNET) of *K. pneumoniae* CG14 and CG15. AcCNET was performed using Pangenome Analysis Toolkit (PATO) and enrichment functions. Label nodes represent *K. pneumoniae* CG14 and CG15 genomes colored according to the clade/capsular-type. Unlabelled nodes represent protein families. Protein families colours represent the -log10 (*p*-value) of the enrichment test. The nodes were arranged using ForceAtlas2 algorithm. Edges are removed to improve the quality of visualisation.

### Plasmidome analysis

We detected 2463 plasmids, 533 of them clustered in 27 plasmid groups (PG) composed by 5 up to 59 plasmids, 184 in PG containing 2-4 plasmids and 333 were singletons. Figures 3 and 4 reflect the diversity of the plasmids within each CG according to the size, the content in replicons and relaxases and predicted mobility. Interestingly, most plasmids were predicted as non-mobilizable (60%; 1477/2463) or mobilizable (32%; 782/2463) based on the absence of conjugative machinery and/or relaxase, and a minority were predicted as conjugative (8%; 204/2463).

**Figure 3.**
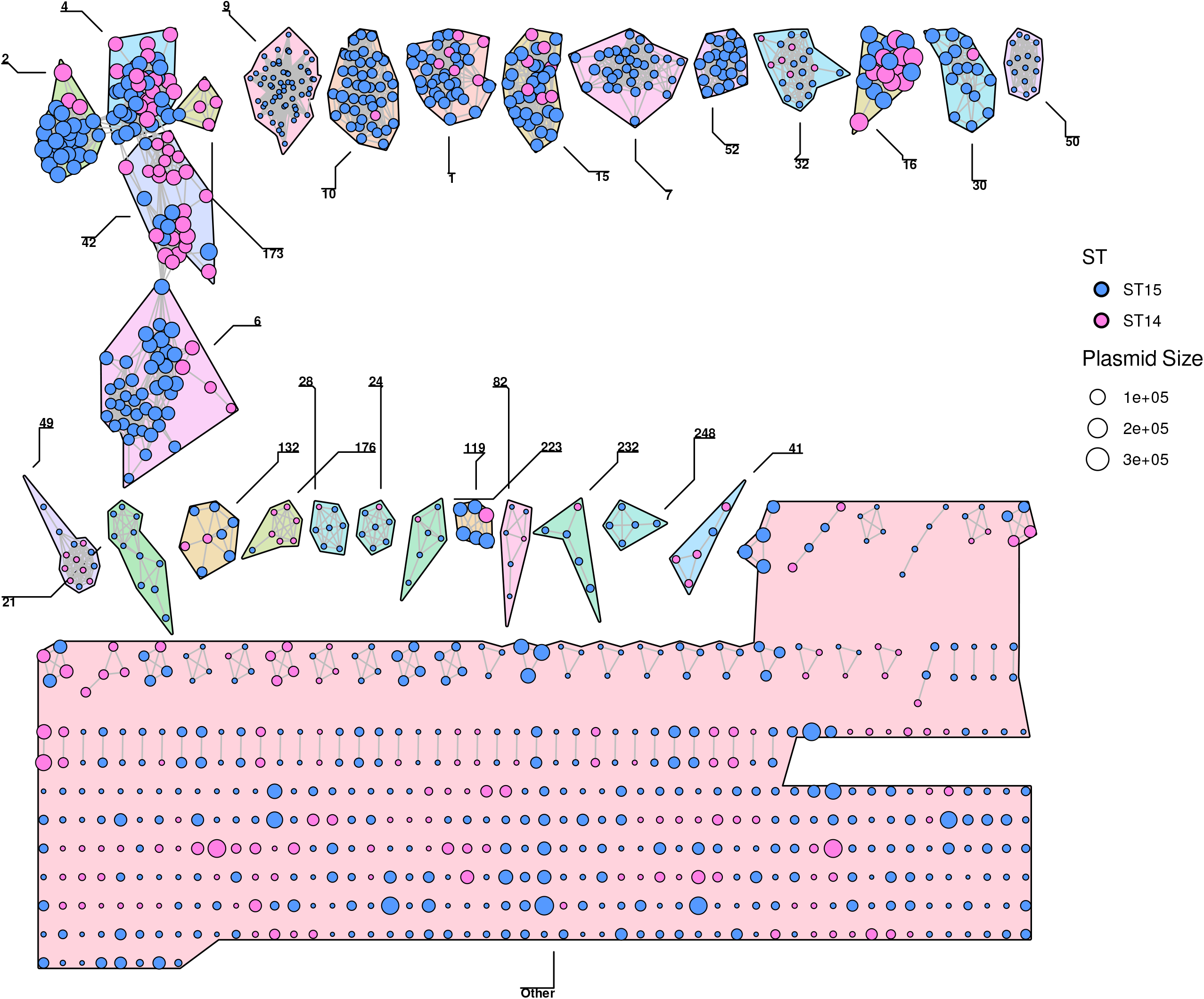
K-nearest neighbour network (K-NNN) of *K. pneumoniae* CG14 and CG15 plasmids. Network was built using the PATO k-nnn function. Each node represents a plasmid and is colored according to the CG. Each plasmid is connected with the 10 best hits if their Jaccard similarity is at least 0.5. Plasmid clusters are defined using the Louvain algorithm over the network structure.

**Figure 4.**
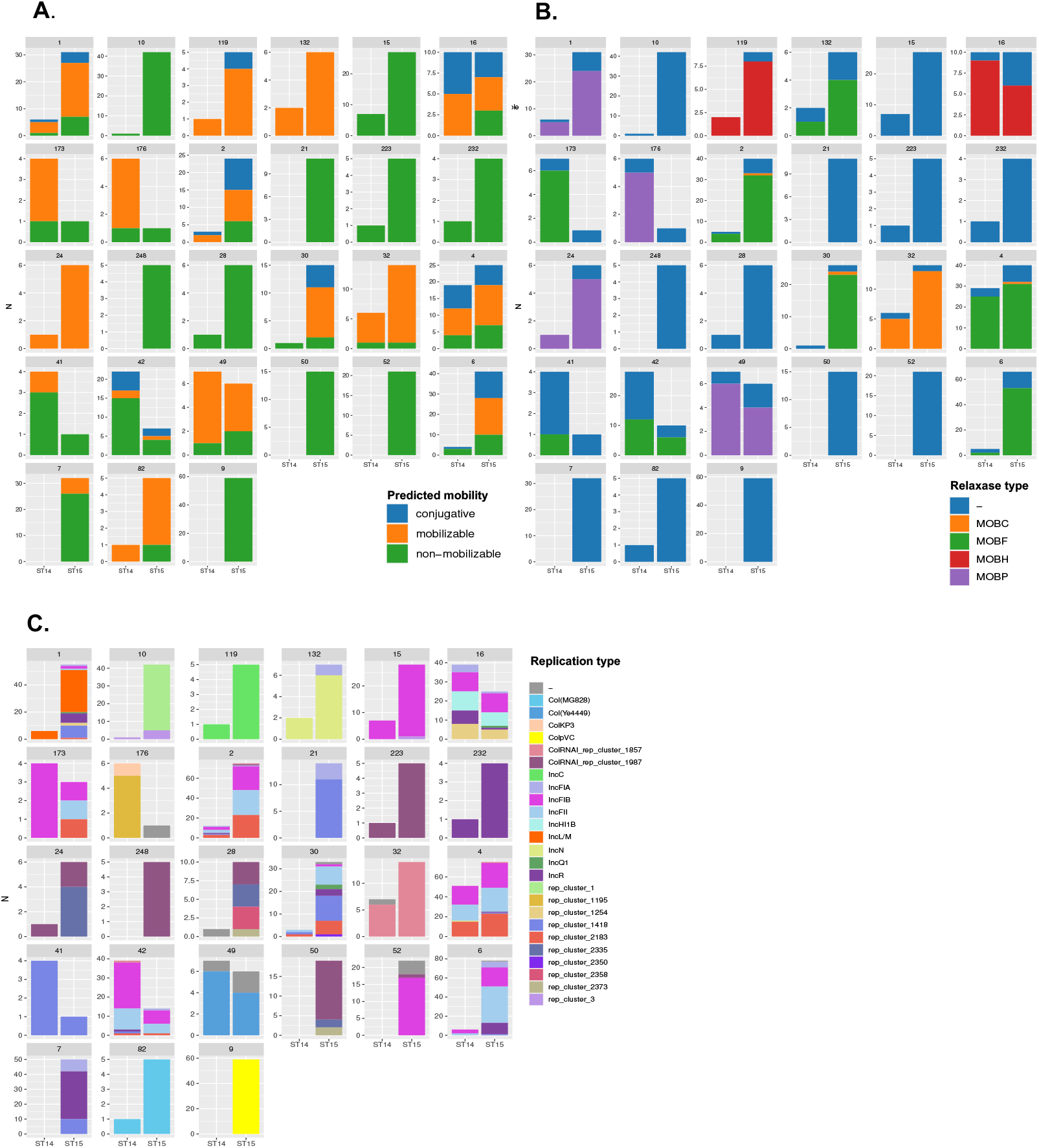
Predicted mobility, replicases and relaxases of each plasmid group (PG). **Panel A**: Distribution of plasmid predicted mobility within each PG in CG15 and CG14. **Panel B**: Distribution of replicases (incompatibility group) within each PG in CG15 and CG14. **Panel C**: Distribution of relaxases (MOB type) within each PG in CG15 and CG14.

The predominant PGs (detected in >10 isolates) carried replicons of F-type (10 PGs), L-type (PG-1), Col-type (PG-9, −32, −49 and −50), and other still undescribed replicon sequences (PG-10, PG-21). IncN (PG-132) or IncC (PG-19) were only sporadically observed, as well as other PG with broad host range such as PG-50. Of note, the F-like PGs (n=10/27, 50-300Kb) differed in the number and type of replicons and relaxases, all with a narrower host range (*Enterobacterales* or *Enterobacteriaceae* family) (**Figure 4, Figure S6**). Six out of 10 F-like PGs had relaxases of the MOBF family. Combinations of FIB(k) (6 PG), three new replicases (5 PG) and FIIk (FII_K1_, FII_K4_ in n=5 PG), R (3 PG), FIA(HI1) (3 PG) and HI1B (1 PG) were observed. Of note, the most abundant PGs were mosaics with regions highly homologous to those of known hybrid plasmids of *Kp* (e.g. pKPN3, pKPN307-D, pKP6402 or pUUH239) (**Table S5**).

A diversity of Col plasmids was also identified (8/27 PGs). Here, the replicon ColpVC (PG-9, 2Kb) of pVCM1 from *Salmonella enterica* was the most abundant (35%; 59/170 isolates), encountered in in CG15 genomes. Other Col replicons were ColRNA_rep1857 (PG-32, 9-12kb), ColRNAI_1987 (2-12kb; PG-50, PG-223 and PG-248), ColYe4449 (PG-49, 6Kb), or ColMG828 (PG-82, 1.6Kb). It is of remark that most new replicases, FIA and Col have been associated with plasmids from *S. enterica* (ColpVC, rep_2183, FIA), *Yersinia enterocolitica* (ColYe4449), *Acinetobacter baumannii* (rep_1254), *Escherichia coli* (4 Col including ColMG828), *Serratia marcescens* (rep_2335) or *Aeromonas hydrophila* (rep_1195) (**Table S5**).

Most PGs (74%, n=20/27) were detected in both CG15 and CG14 with variable frequency, only a few of them (PG-4, −16, −49, −173; F-type or Col) being equally distributed in both CGs. Some PGs were associated with CG14 and especially in subclade KL2 (PG-41, −42, −176; F-type or Col). Most PGs were present in CG15 and distributed in different subclades. Of remark, seven PGs were exclusively found in CG15 (PG-7, −9, −10, −21, −50-, 52, −248) corresponding to multireplicon F (n=3), Col (n=3) or FIB (n=1), all lacking MOB. Some PGs were abundant in KL24 (PG-2, −4, −28, −30, −248) or KL112 (PG-6, −7, −21, −50, −52) or both (PG-1, −9, −10) (**Figure S7**).

### Distribution of antibiotic resistance and metal tolerance genes

ARGs were located on mainly plasmids from IncF, IncC, IncL, IncN, and Col type (**Figure S8**). Plasmid sequence analyses reveal diverse modes of ESBL and carbapenemase gene spread among *Kp*. The *bla*_CTX-M-15_ gene was mostly associated with *bla*_TEM-1A_, *dfrA14*, *aph(6)-Id*, and *aph(3’)Ib* on a diversity of plasmid backgrounds (mostly F-type, L and C), probably reflecting the promiscuity of a widespread transposable unit (31). *bla*_NDM-1_ appears also in different plasmid entities (IncC, IncN, IncHI1B). On the contrary, the *bla*_OXA-48_ was only located on IncL plasmids related to pOXA-48 (Genbank acc. number JN62686), *bla*_OXA-232_ by the new Col (rep_1195), *bla*_TEM-24_ or *bla*_CMY_ was associated with IncC and *bla*_SHV-12_ or *bla*_KPC-2_ with IncN. Of note, *bla*_VIM_ and *bla*_KPC-3_ were rarely detected.

It is of interest to note that certain predominant plasmid types carry diverse ESBL or carbapenemase genes, namely IncC (*bla*_CTX-M-15_, *bla*_NDM-1_, *bla*_SHV-5_, *bla*_CMY_, *bla*_TEM-24_), IncN (*bla*_KPC-2_ *bla*_NDM-1_ or *bla*_SHV-12_) or the F-HI1B multireplicon plasmid (PG16) carried the highest number and diversity of ARG including *bla*_NDM-1_ or *bla*_CTX-M-15_, and showed regions identical to reference NDM-1-encoding plasmids as pPMK1-NDM and pNDM-MAR (Genbank acc. numbers CP008933 and JN420336, respectively).

Metal tolerance genes (MTGs) were also enriched in multireplicon and multidrug resistance plasmids of the incompatibility groups F (n=7 PGs), L and C (mercury operons in PG-7, copper (*pco*), silver (*sil*) and arsenic (*ars*) operons in F plasmids) (**Figure 1**).

## DISCUSSION

The phylogenomic analysis of available CG15 and CG14 genomes of this study reveals that these two globally distributed *Kp* clonal groups have an independent evolutionary trajectory. The recent nomenclature proposed to different *K. pneumoniae* sublineages is in line with these findings (17). The deep branching structure of their core genome shows a diversification that involves capsular switches and further expansion of the dominant clades (KL2 in CG14; KL24/KL112 in CG15). Time-scale inferences suggest the emergence of KL2- and KL16-CG14 subclades predate that of main CG15 subclades, whose seem to have emerged contemporary to other globally distributed lineages around the 1990’s (ST307, ST147, ST101, ST258) (7–10).

Though our sample is clearly dominated by genomes from European countries, CG14 and CG15 subclades seem to be globally distributed (11, 32), similarly to other subclades within CG147, CG258, CG307 or CG101. This suggests their specialisation through acquisition of adaptive traits that seem to favour colonisation, spread and persistence in shifting conditions (7–10). Recombination events involving the capsule locus could have allowed the adaptation to human immunity and consequent selection and amplification of certain *Kp* subpopulations (33–35), similarly to what has been described in *Streptococcus pneumoniae* or *Neisseria meningitidis* (36, 37). In fact, the recognition of these K-types as suitable evolutionary markers led to their use in diagnostics for categorising *Kp* isolates (including CG15 and CG14) in hospital surveillance and infection control programs using Fourier Transform Infrared spectroscopy (30, 38, 39).

Besides the capsule, the acquisition of resistance to fluoroquinolones by most CG15 through specific mutations in GyrA and ParC also mimics what has been reported for other contemporary high-risk clones of *Kp*, *Escherichia coli*, and other species (7, 8, 40–42). In fact, fluoroquinolone resistance seems to be an advantage for the fitness of high-risk clones of different species (40, 41). Fluoroquinolone resistant clones predominate in the elderly and people with previous exposure to healthcare centers that facilitates the acquisition of other antibiotic resistances (43, 44).

This study presents one of the few available comprehensive studies on the composition of the entire *Kp* plasmidome. Such analysis reveals the overrepresentation of F-type and Col plasmids which are more diverse than previously recognized for a given CG. The occurrence of such plasmids in other MDR *Kp* populations is unknown but the high similarity to plasmid backbones described in other CGs (e.g. mosaic pKPN-3-like plasmids) suggests their wide distribution and their role in the acquisition and spread of contemporaneous ESBL and carbapenemase genes (*bla*_CTX-M-15_, *bla*_KPC_, *bla*_NDM-1_) (45–49). It is of note the occurrence of plasmids harbouring “novel” plasmid replicons that are unrecognisable by current plasmid classification schemes (23, 50–52) (**Table S5**). The high occurrence of non-conjugative/non-mobilizable plasmids is not surprising (50), and implies either the high recombination of the mobile genetic elements in high-risk clones, with frequent deletion and acquisition events and/or the need of helper entities (e.g. conjugative plasmids, ICEs) for their mobilisation (53–55). Carriage of genes encoding adaptive functions (antimicrobial resistance, metal tolerance, thermoresistance, amongst others) reflects a history of exposure to disparate selection pressures (54). The high degree of mosaicism observed, especially on F-type plasmids is in agreement with previous observations, and is suggestive of the impact of recombination in the dissemination of ARGs (56, 57). The high level of recombination between highly similar plasmids and replicons, and the narrow host range of most plasmid groups (at the genus-*Klebsiella* and family-*Enterobacteriaceae* level) suggests a degree of specialization that results from the long-term co-evolution of these plasmid types in this genetic background (**Figure S6**) (55).

It is also of note the high heterogeneity of small plasmids (essentially ColE1-type) long evolving in Enterobacterales, some of them particularly abundant and linked to CG15 (ColpVC, ColRNAI_rep1987). Col plasmids have been implicated in extraintestinal virulence and/or gastrointestinal colonisation in *E. coli* (53, 58). Their occurrence in *Kp* and their function are largely unknown but they contribute to the dissemination of antimicrobial resistance determinants and other host adaptive factors (cell metabolism, virulence, defence from phages or heavy metal resistance) as well as to plasmidome plasticity and evolution through events mediated by common transposases (*IS26, IS4321*, Tn*3*-like) (21, 53, 59, 60)(this study).

The analysis of the accessory genome of *Kp* CG14 and CG15 reveals the presence of ARGs, long associated with selection and diversification. Second, the acquisition of ARGs and/or MTGs by multiple plasmid backgrounds contributed to the diversification and expansion of CG15, and particularly the KL112 clade. In fact, *bla*_CTX-M-15_ is the most frequently acquired β-lactamase encoding gene (38% in ST14 *versus* 68% in ST15), increasing in prevalence from ST14-KL2 (45%), ST15-KL24 (62%) and KL112 (92%). The repeated acquisition of the multidrug resistance cassette containing *bla*_CTX-M-15_ by multiple F-type plasmids might have contributed for this expansion, as reported for *E. coli* ST131 (61). Nonetheless, the presence of genes encoding ESBL, AmpC and carbapenemases or MTGs by promiscuous and multidrug resistance plasmids (IncC, IncN, IncL/M) might have also contributed for the selection of the ST15-KL112 clade, representing an increasing clinical challenge (18, 62).

## CONCLUSIONS

This study demonstrates an independent evolutionary trajectory for CG15 and CG14, marked by expansion of subclades with particular capsular types (CG14-KL2, CG15-KL24/KL112), GyrA and ParC mutations (within CG15) and ARGs in a high diversity of mosaic plasmids. A comprehensive analysis of the plasmidome revealed the contribution of a turbulent flux of plasmids, particularly pervasive F-type and Col plasmids, and adaptive traits (ARGs, MTGs, VFs) to the pangenome, which reflect the exposure to different selective pressures of MDR *Enterobacteriaceae*. Finally, this study further highlights the relevance of plasmid data for the understanding of ARGs spread and CG evolution, and the diverse trajectories of ARGs in clinical settings (one plasmid/multiple lineages, multiple plasmids/multiple lineages, and multiple plasmids/one lineage).

## MATERIAL AND METHODS

### Bacterial Isolates and Sequencing

Seven MDR ST15 (3 K24, 1 KL39, 1 KL112, 1 KL19, 1 KL110) and two ST14 (1 KL2, 1 KL16) MDR isolates representing main CG15 and CG14 circulating in Portugal and elsewhere between 2003 and 2013 were selected for whole-genome sequencing (WGS) (27–30) (**Table S3**). Bacterial genomic DNA was extracted using QIAmp DNA Mini Kit (Quiagen), and DNA concentration and purity measured using Qubit Fluorometer (Life Technologies) and Nanodrop 2000 (Thermo Scientific), respectively. DNA libraries were prepared using the Nextera XT kit (Illumina, San Diego, CA, USA) and 2 x 300 bp paired-end sequence reads with mean coverage of 100x were generated on Miseq platform (Illumina, San Diego, CA, USA). *De novo* assembly was performed with SPAdes v3.9.0 using *k*-mers of 101, 111, 121 and 127 (63), and the quality of the assemblies evaluated using QUAST (64). The whole genome shotgun project was deposited in DDBJ/EMBL/GenBank under the BioProject accession number PRJNA408270, and assembly statistics concerning the nine isolates sequenced in this study are available on **Table S3**.

### Public available genomes and Global dataset

A total dataset of 481 publicly available CG14 (n=110; 1980-2018) and CG15 (n=371; 1980-2017) *Kp* genomes from NCBI RefSeq (November 2019) were downloaded, provided they complied with the control quality criteria defined [genome size (4.9-6.2Mb) and %GC (56-58%) matching with *Kp*, and less than >1000 contigs or N50 >20K] (**Table S6, Figure S9**). From these, all the genomes without available information concerning the isolation year (n=20) were excluded. Furthermore, 226 genomes (30 CG14 and 196 CG15) showing <21 Single Nucleotide Polymorphisms (SNPs) in the core-genome alignment obtained with Roary were discarded since they were considered epidemiologically related (11). Furthermore, to evaluate the strength of the temporal signal of our data set and to depict problematic or erroneous sequences (‘outliers’), we conducted a linear regression analysis of the root-to-tip genetic distances as a function of the sample collection year (**Figure S2**), using TempEst v1.5.3 (http://tree.bio.ed.ac.uk/software/tempest/), after which we discarded 9 additional genomes, resulting in 226 deduplicated high quality genomes (**Table S1**).

Our global dataset (n=235 genomes) included 65 CG14 (63 ST14 and 2 SLV) and 170 CG15 (162 ST15 and 8 SLV) originated mainly from Europe (55%), Western and South-eastern Asia (16%) and North America (7%) (**Figure S4**). The vast majority were recovered from humans (97%, 75% in the context of infection) and carried genes encoding for extended-spectrum β-lactamases, qAmpCs and/or carbapenemases (81%), reflecting a strong sampling bias towards multidrug resistant (MDR) strains from the nosocomial setting. All the epidemiological and genomic data concerning the final dataset of genomes are shown in **Table S1**.

### Core Genome Phylogenetic Analysis

To conduct the phylogenetic analyses, the 235 genomes from our final dataset were previously annotated using Prokka 1.12 (65) and a core-genome alignment was constructed with Roary v3.12 (66) using a blastP identity cut-off of 90% and core genes defined as those being present in more than 90% of the isolates, resulting in a total of 4,420 core genes. Afterwards, 35,783 single-nucleotide variants (SNVs) were extracted from the core-genome alignment with SNP-sites (67) and used to construct a maximum-likelihood phylogenetic tree with IQ-TREE v1.6.11 (model GTR+F+ASC+R4) rooted with two closely related outgroup genomes (15): *Kp* ST540 Kpn0019 (accession number: SRR2098710) and a *Kp* ST101 Kp_Goe_33208 (GCF_001902435.1) (**Figure 1**).

After inspection of the global tree, we have generated a recombination-free alignment using Gubbins v1.12 (68) and evaluated the strength of the temporal sign with TempEST for the whole dataset (R^2^=0.04733, **Figure S2A**) and for the two CGs independently (R^2^=0.04733, **Figure S2BC**), and decided to perform the temporal phylogeny analysis independently for each CG with BEAST v2.6.2 (run with a Markov chain Monte Carlo length of 1×10^9^, sampling every 5×10^3^ steps) (69). We used model parameters that had the best fit: GTR substitution model, lognormal relaxed clock and constant population size. Parameter estimates were computed using Tracer v1.7.1, and a maximum clade credibility tree was obtained with TreeAnnotator v2.6.0. and visualised in FigTree v1.4.4 (69).

### MLST/cgMLST and genomic analyses of surface polysaccharides, antimicrobial resistance, virulence and heavy metal tolerance

MLST and cgMLST was performed using the Institut Pasteur Klebsiella MLST database (https://bigsdb.pasteur.fr/klebsiella/) (6, 17). Genes associated with antimicrobial resistance and virulence [siderophores (yersianabactin, aerobactin, colibactin, salmochelin), regulators of mucoid phenotype (*rmpA* and *rmpA2*)] were searched using Kleborate v1.0 (https://github.com/katholt/Kleborate/wiki) (12). Characterization of the surface polysaccharides, capsular locus (KL) and LPS O-antigen, was performed through Kaptive, integrated in Kleborate (12, 70). Additional virulence factors, such as iron uptake systems (*kfu* operon), bacteriocins (microcin cluster), and adhesins (*mrk* cluster), as well as heavy metal tolerance genes [arsenic (*ars* operon), copper (*pco* cluster), silver/copper (*sil* cluster), tellurite (*ter* operon) and mercury (mer operon)] were searched using BIGSdb (https://bigsdb.pasteur.fr/klebsiella/), whereas the *kpi* chaperone-usher pili system was searched using BLAST (26). Statistical analyses to check the association of the different categorical variables within the clades defined were calculated using the *χ^2^* test (*P* values of <0.05 were considered statistically significant).

### Accessory Genome Analysis

The accessory genome was defined as all genes present in less than 80% of the genomes and was obtained using PATO with default parameters (protein families: 80% identity, 80% coverage) (25), followed by a detailed study of the protein enrichment in each of the CGs (**Table S4**). The table shows the genes enriched in some of the CGs with a p-value lower than 0.001 and present in more than 75% of the genomes of the CG. To illustrate the accessory network, we use Gephi for the rearrangement and then we load the layout in R to plot the network.

### Plasmid identification and analysis

A preliminary analysis of the plasmid content of CG15 was obtained by searching the diversity of plasmid replicons on the genomes using MOB-suite (52). To analyse the plasmidome a population structure of the plasmids was created using a similarity network based on the presence/absence data. Using the presence/absence matrix, we calculated the similarity among the plasmids using Jaccard distance. Then we create a k-nearest neighbors network (K-NNN) with PATO with parameters 10 neighbors, allowing reciprocal connections and with a similarity threshold of 0.5 (Jaccard Distance) (**Table S5**). Finally, we incorporate the plasmid typing information (predicted mobility, replicases and relaxases) to the network (71).

## Funding

This work received financial support from the European Union (FEDER funds POCI/01/0145/FEDER/007728) and National Funds (FCT/MEC, Fundação para a Ciência e Tecnologia and Ministério da Educação e Ciência) under the Partnership Agreement PT2020 UID/MULTI/04378/2013. Carla Rodrigues was supported by a fellowship from FCT through POCH (grant number SFRH/BD/84341/2012) and a FEMS Research Grant (FEMS-RG-2014-0089). Ângela Novais is supported by national funds through FCT, I.P., in through Scientific Employment Stimulus Program [2021.02252.CEECIND/CP1662/CT0009]. TMC’s lab is supported by the European Commission (grant MISTAR AC21/2 00041) and the Instituto de Salud Carlos III (ISCIII) of Spain, cofinanced by the European Development Regional Fund (A Way to Achieve Europe program; Spanish Network for Research in Infectious Diseases grants PI18/1942; PI21/02027; CIBER CB21/13/00084) and the Fundación Ramón Areces (BioMetasep).

## Competing interests

The authors have declared that no competing interests exist.

## Notes

### Competing Interest Statement

The authors have declared no competing interest.

