## Supplementary material for "Phylogenomics of globally spread Clonal Groups 14 and 15 of *Klebsiella pneumoniae*": Figure S1 to Figure S9

**Ângela Novais**

***Co-corresponding author**

**Teresa M. Coque**

***Contents***

**Figure S1.** Boxplot representing pairwise SNP distances among all the *K. pneumoniae* CG15 and CG14 genomes (A), within CG14 and CG15 clades (B) and within the different main subclades (C).

**Figure S2.** Temporal signal in all the *K. pneumoniae* CG15 and CG14 genomes (A), in CG14 genomes (B) and amongst CG15 genomes (5).

**Figure S3.** Time-scaled phylogeny of *K. pneumoniae* CG14 genomes and their antimicrobial resistance determinants.

**Figure S4.** Worldwide distribution of the main subclades identified within *K. pneumoniae* CG14 and CG15.

**Figure S5.** Time-scaled phylogeny of *K. pneumoniae* CG15 genomes and their antimicrobial resistance determinants.

**Figure S6**. K-nearest neighbour network (K-NNN) of *K. pneumoniae* CG14 and CG15 plasmids coloured according to the host-range.

**Figure S7**. Distribution of the different plasmid groups identified in main CG14 and CG15 *K. pneumoniae* subclades.

**Figure S8.** Heatmap of the frequency of different antimicrobial resistance genes in the plasmid groups identified within *K. pneumoniae* CG14 and CG15 genomes.

**Figure S9.** Phylogenetic structure of the initial dataset analyzed in the study.

***Please see separate Excel file for Table S1 to S6.***

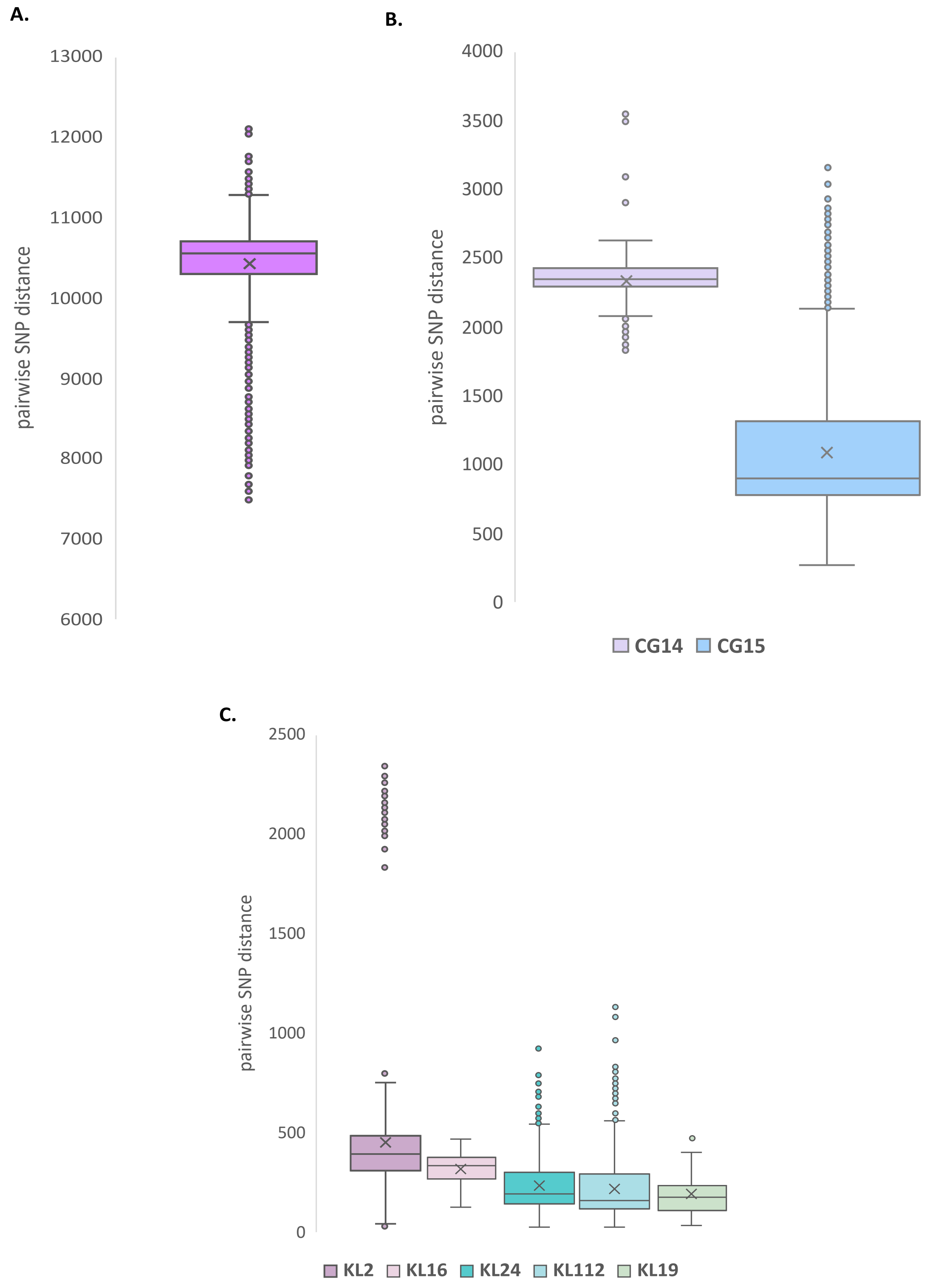


**Figure S1.** Boxplot representing pairwise SNP distances among all the *K. pneumoniae* CG15 and CG14 genomes (A), within CG14 and CG15 clades (B) and within the different main subclades (C).

Box represents interquartile range, whiskers represent the least and greatest values excluding the outliers, solid grey line represents the median and the x represents the mean.

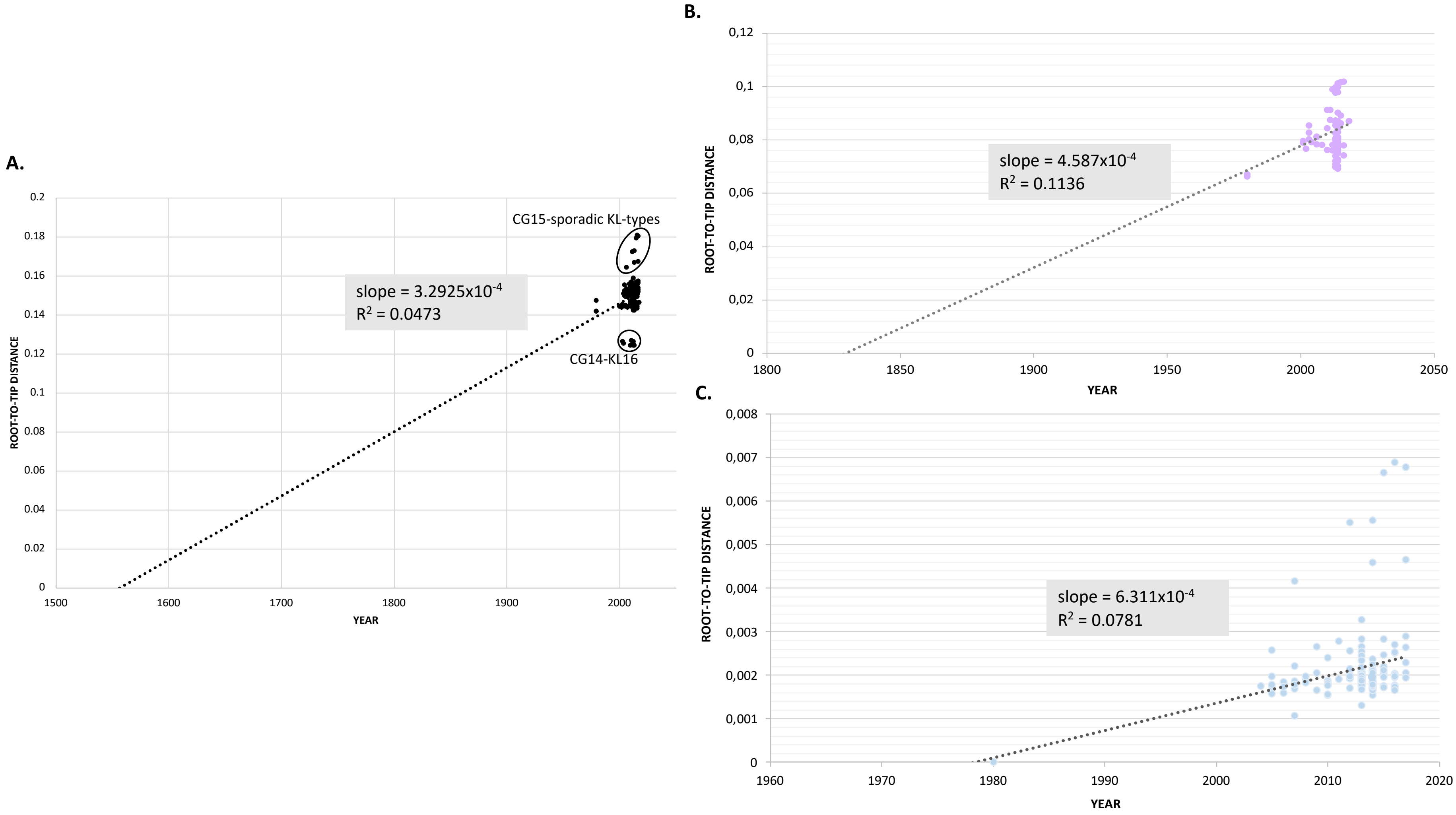


**Figure S2.** Temporal signal in all the *K. pneumoniae* CG15 and CG14 genomes (A), in CG14 genomes (B) and amongst CG15 genomes (C). Linear regression between year of isolation vs root-to-tip distance from the maximum likelihood phylogeny was calculated using TempEst.

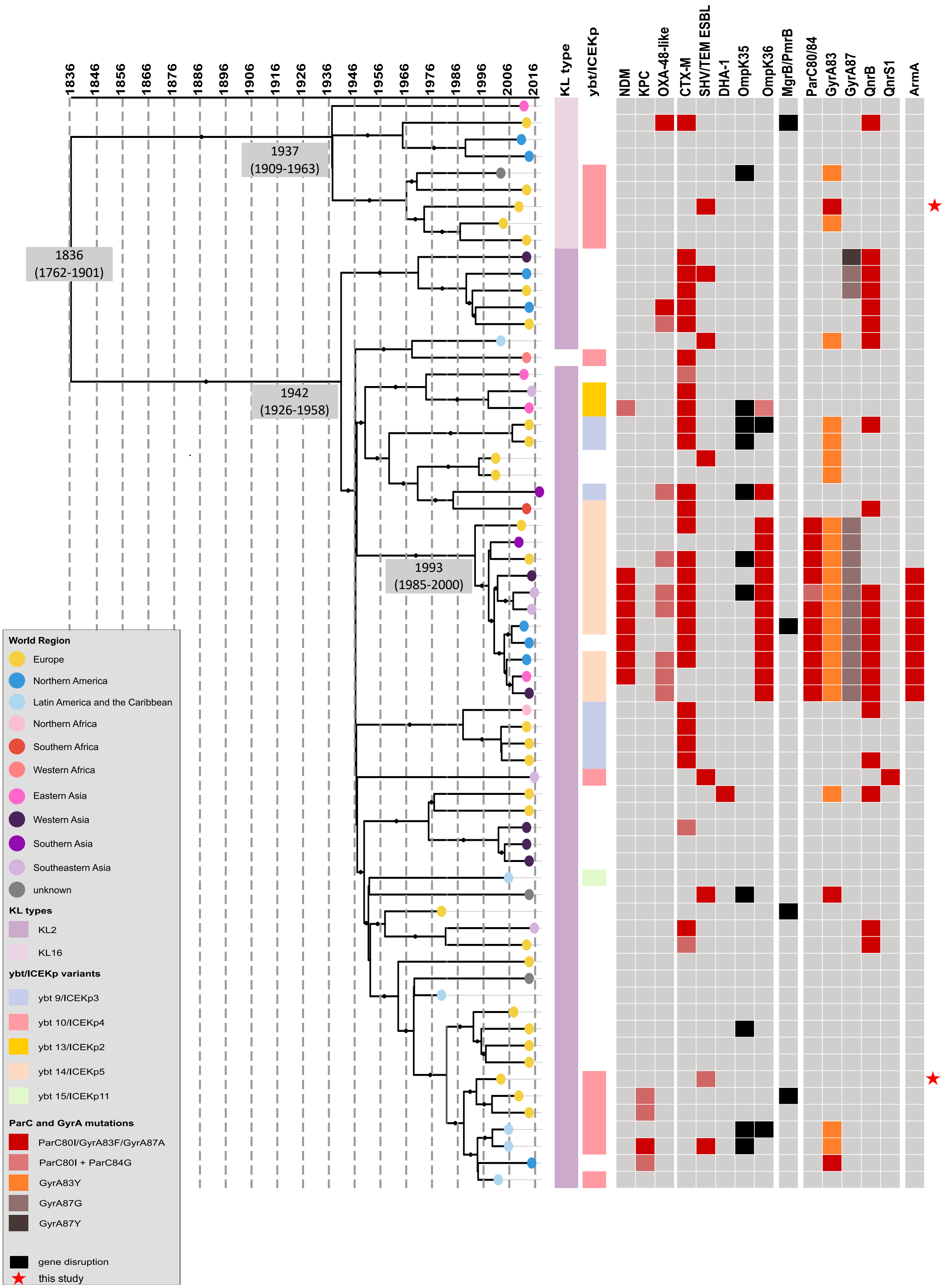


**Figure S3.** Time-scaled phylogeny of *K. pneumoniae* CG14 genomes and their antimicrobial resistance determinants.

Phylogeny obtained with BEAST. The two main branches correspond to the two main clades harboring different capsular-types (KL). Black dots on main nodes indicate ≥95% posterior probability. Tree tips are colored according to the world region of isolation. KL-type and the yersiniabactin-carrying ICE*Kp* elements are colored in accordance with their variants (see key). The presence of different antimicrobial resistance determinants is indicated.

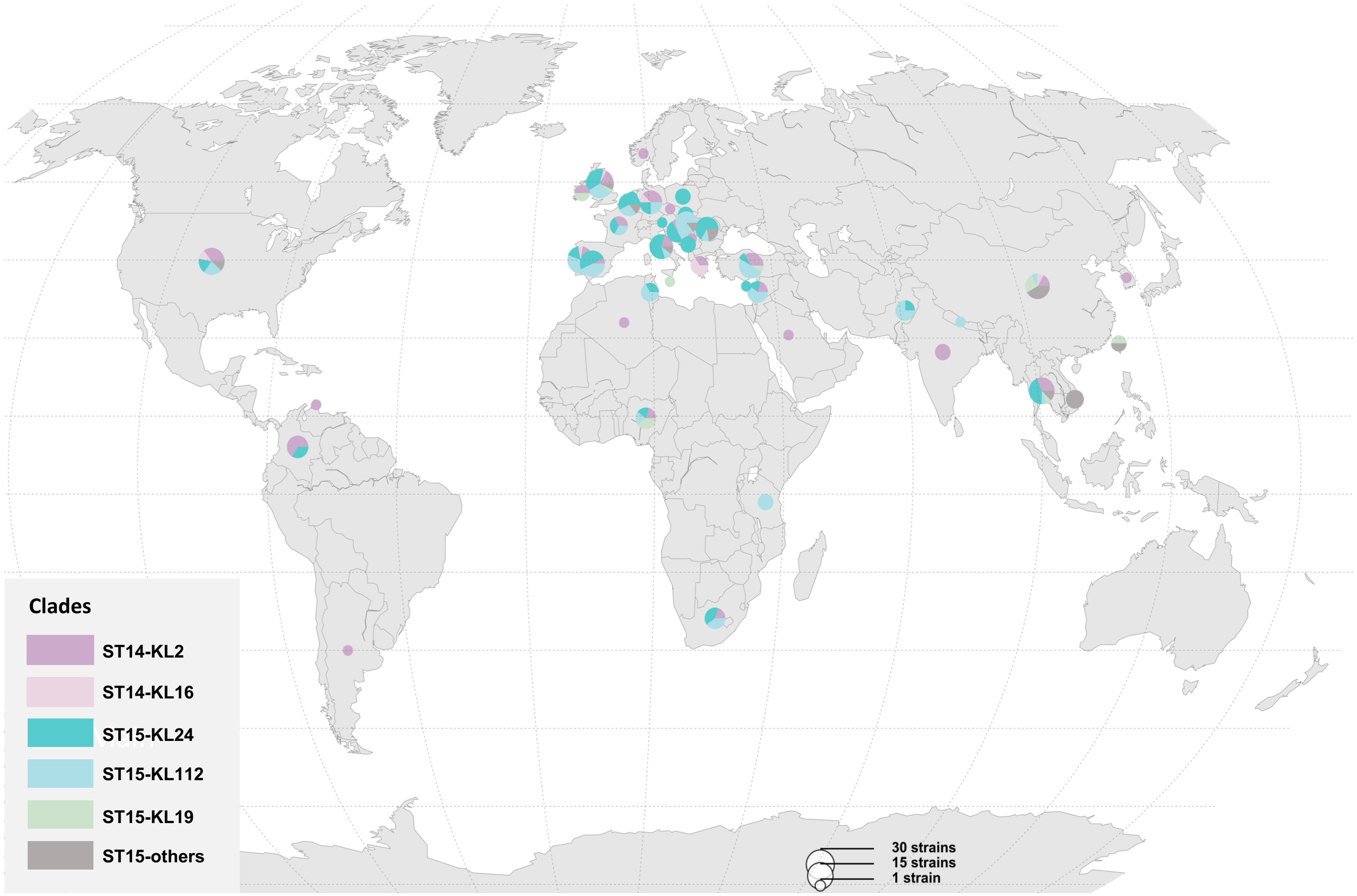


**Figure S4.** Worldwide distribution of the main subclades identified within *K. pneumoniae* CG14 and CG15 genomes included in this study.

The pie charts represent the frequency of each clade in each country (see legend).

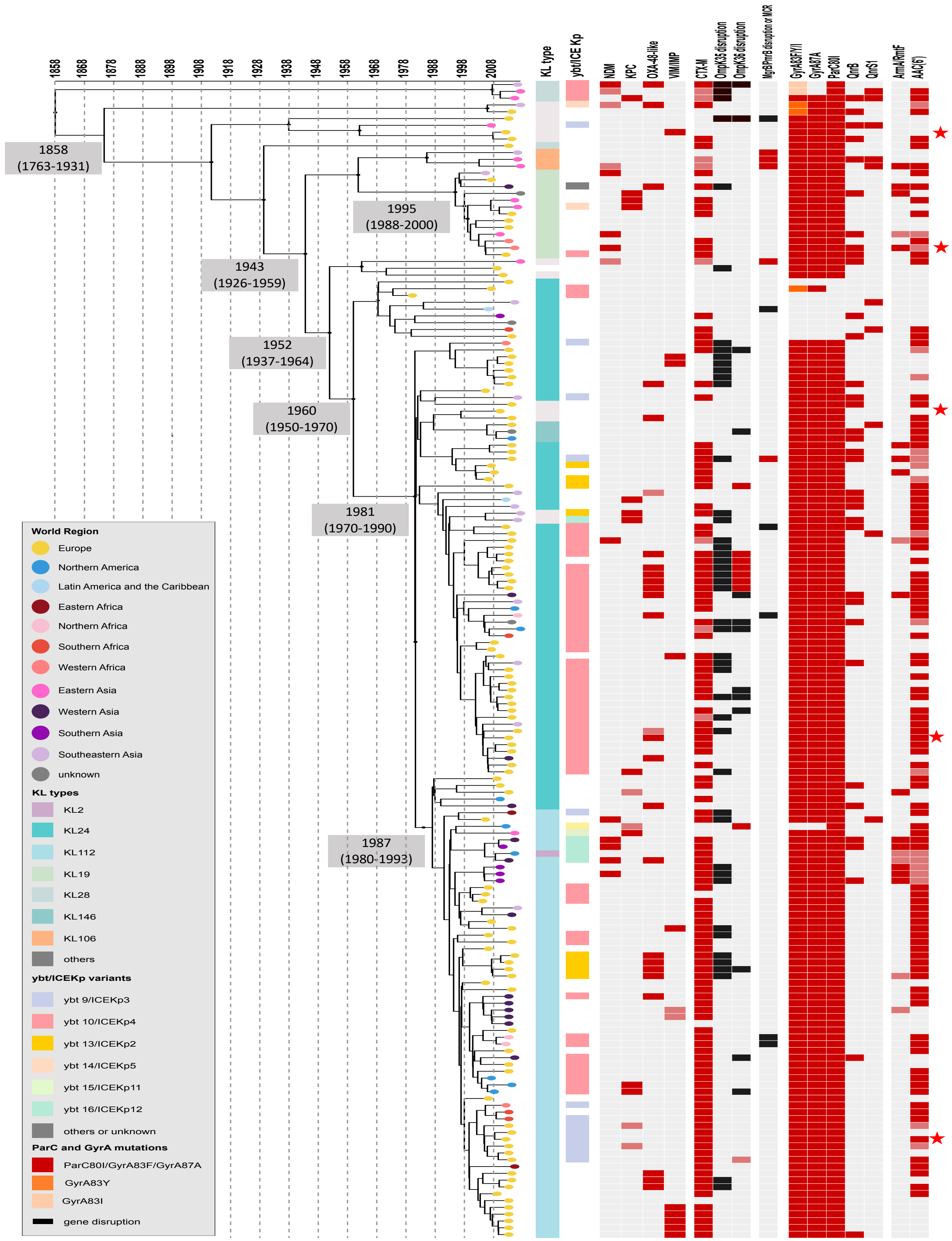


**Figure S5.** Time-scaled phylogeny of *K. pneumoniae* CG15 genomes and their antimicrobial resistance determinants.

Phylogeny obtained with BEAST. Black dots on main nodes indicate ≥95% posterior probability. Tree tips are colored according to the world region of isolation. KL-type and the yersiniabactin-carrying ICE*Kp* elements are colored in accordance with their variants (see key). The presence of different antimicrobial resistance determinants is indicated.

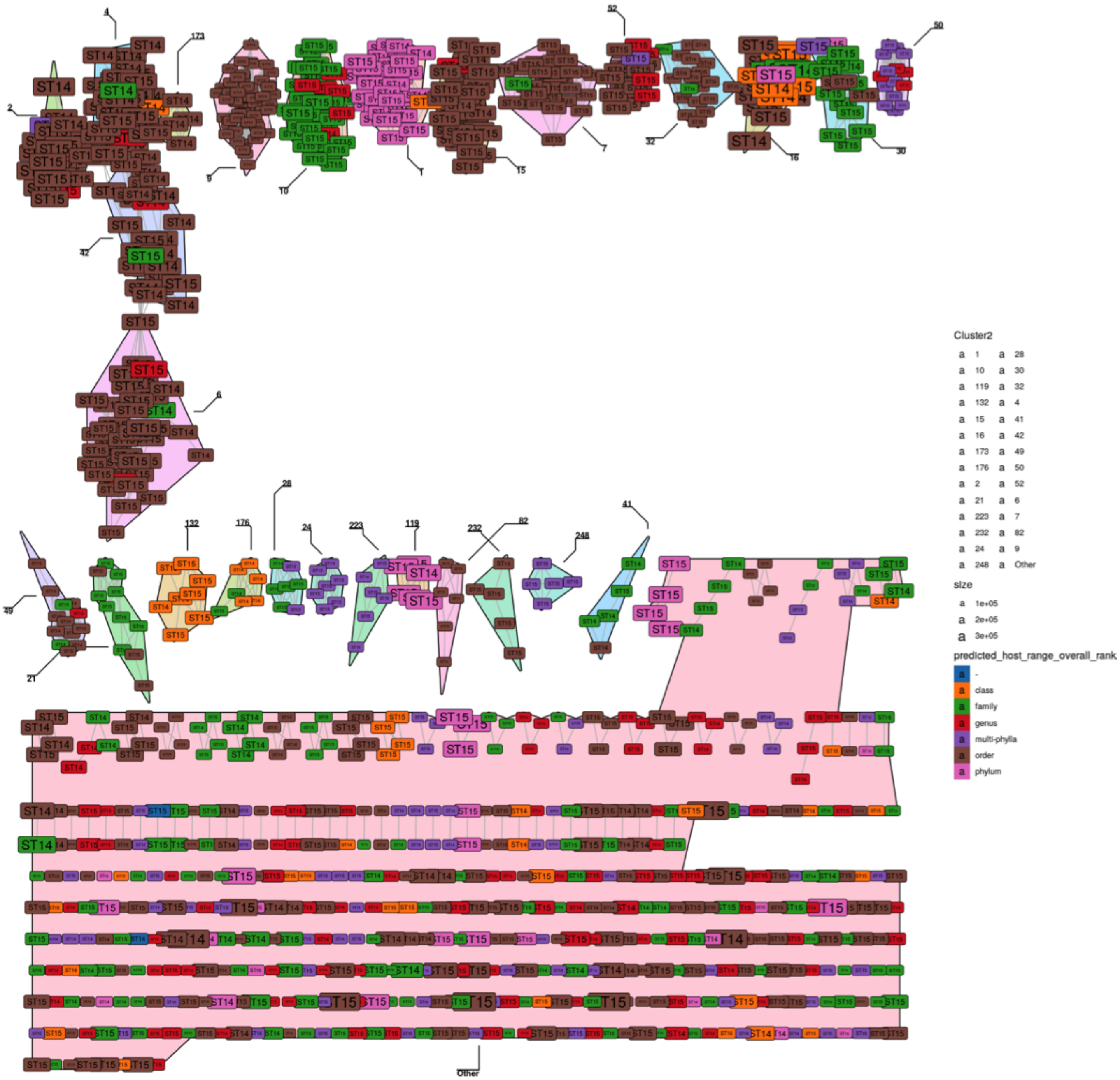


**Figure S6**. K-nearest neighbour network (K-NNN) of *K. pneumoniae* CG14 and CG15 plasmids coloured according to the host-range.

Network was built using the PATO k-nnn function. Each node represents a plasmid and is colored according to the host-range. Each plasmid is connected with the 10 best hits if their Jaccard similarity is at least 0.5. Plasmid clusters are defined using the Louvain algorithm over the network structure.

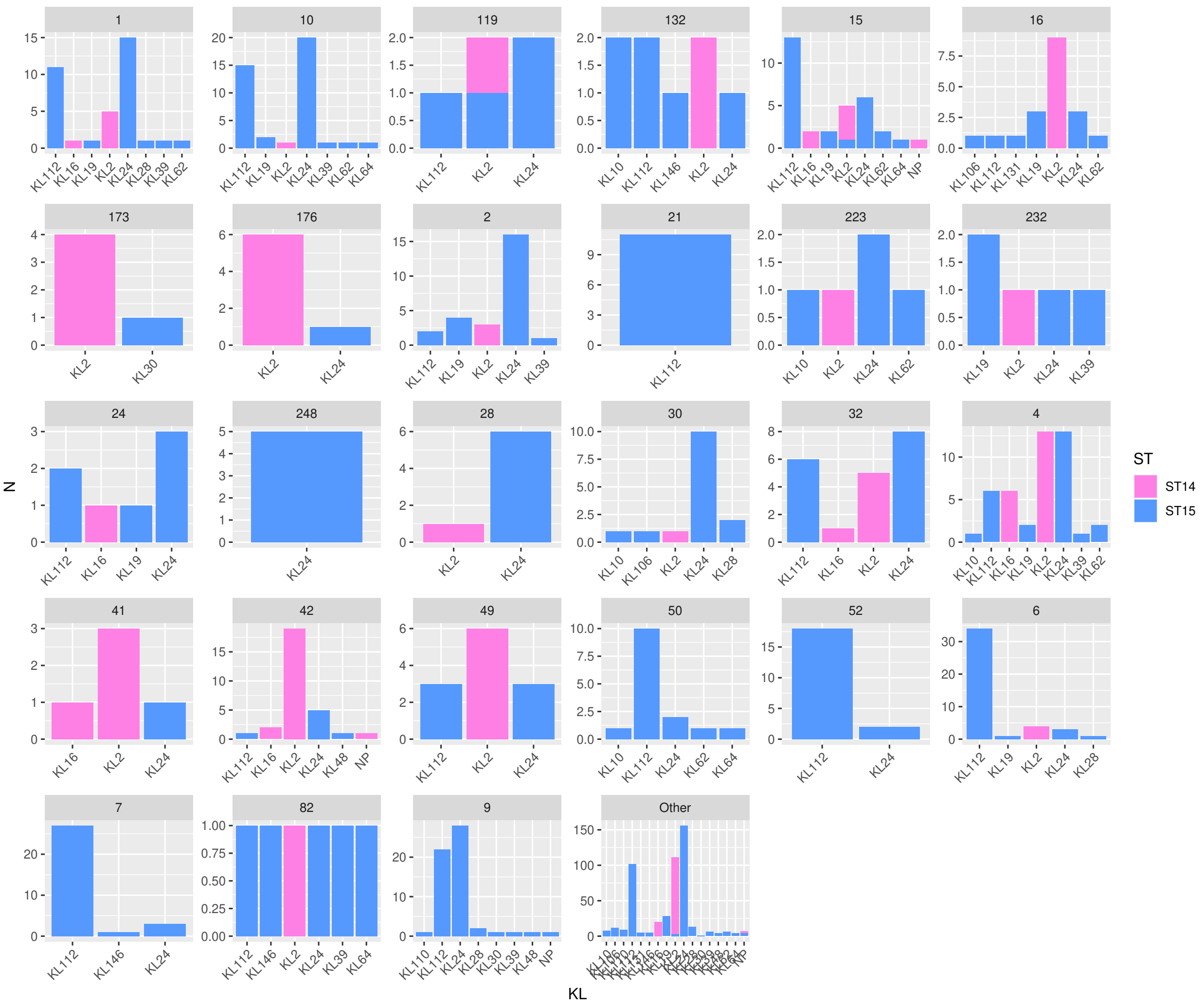


**Figure S7**. Distribution of the different plasmid groups identified in main CG14 and CG15 *K. pneumoniae* subclades.

N, number of genomes, KL, capsular-type

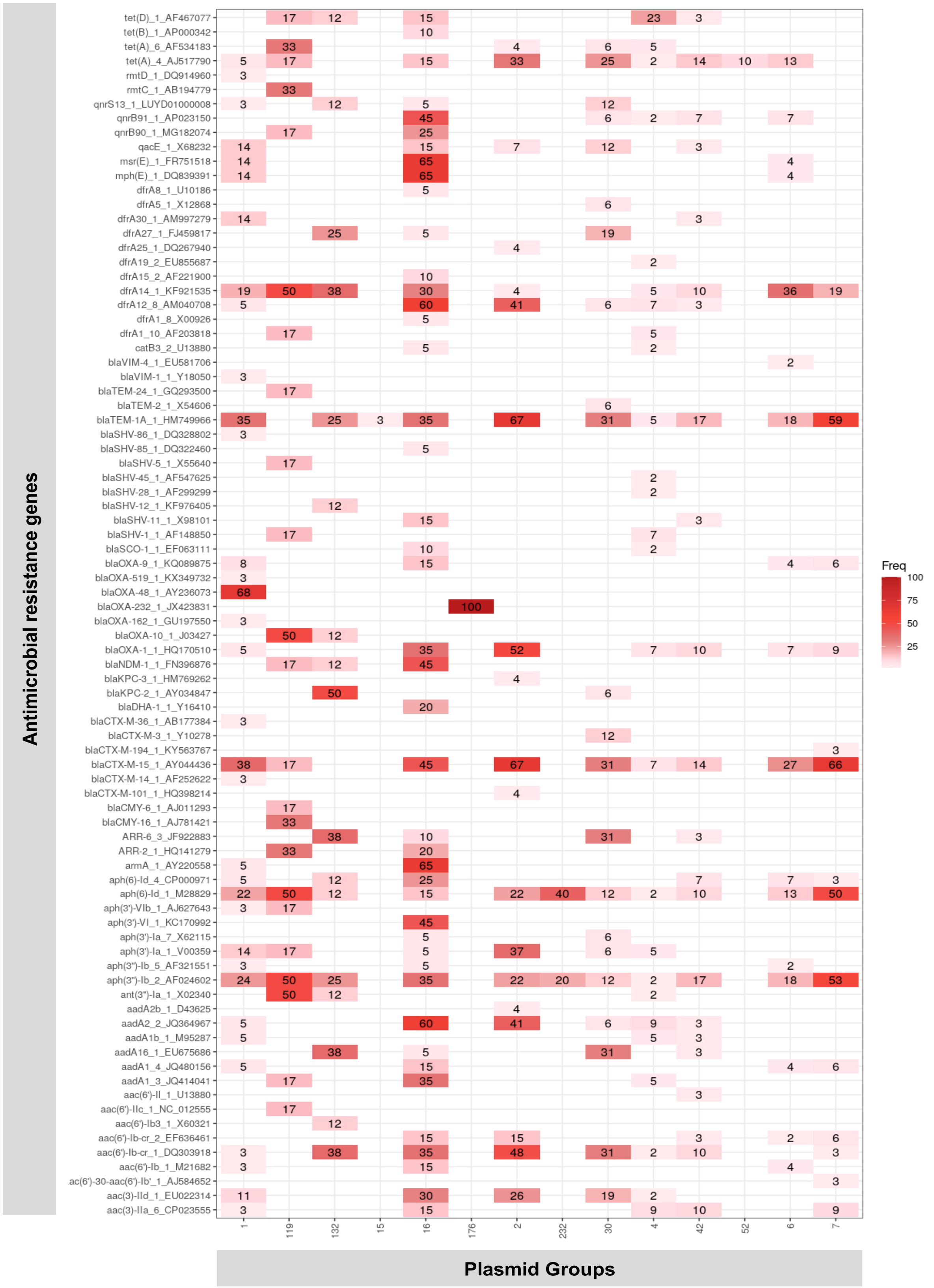


**Figure S8.** Heatmap of the frequency of different antimicrobial resistance genes in the plasmid groups identified within *K. pneumoniae* CG14 and CG15 genomes.

The absolute number of AMR genes identified in each plasmid group is indicated in each square.

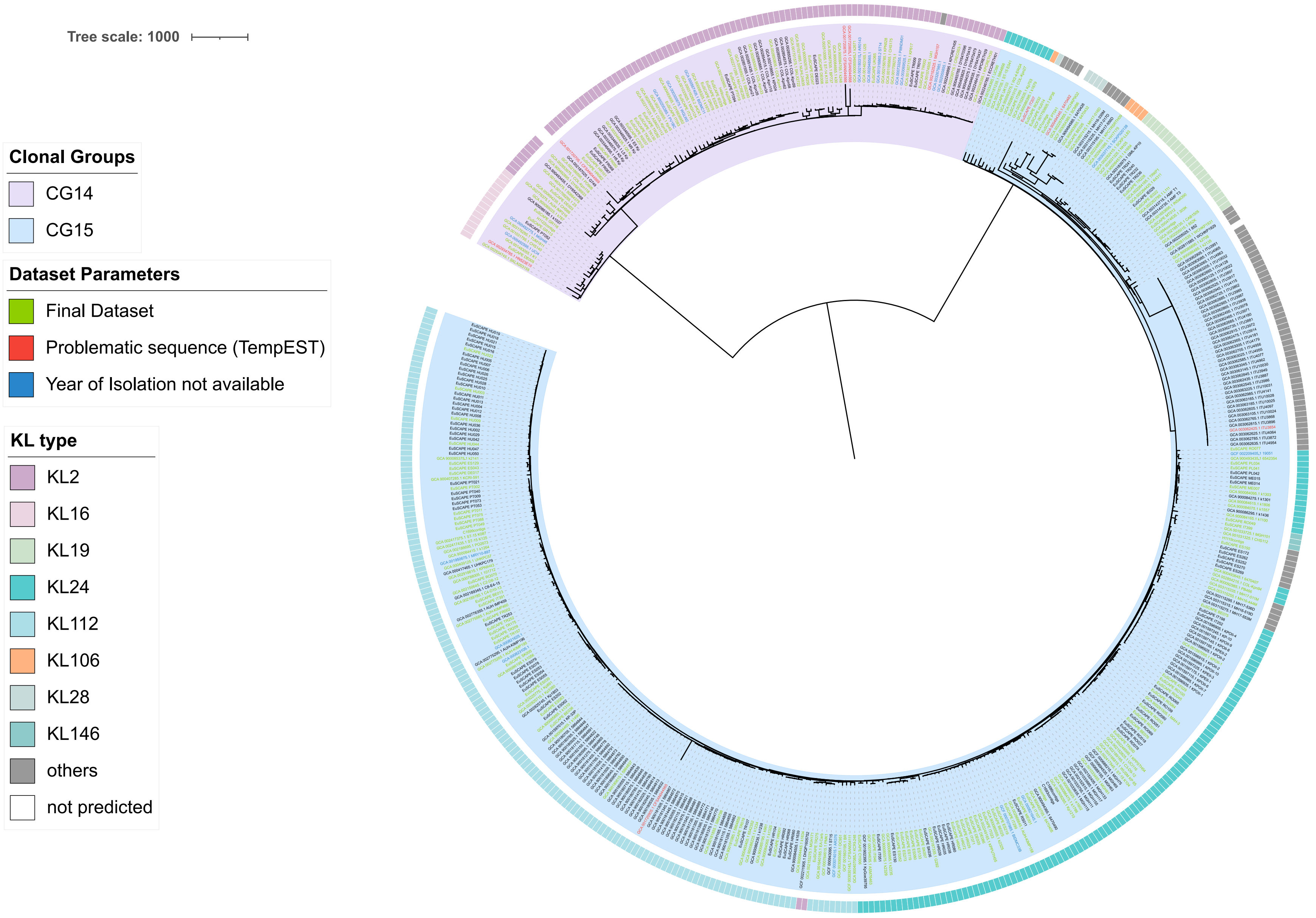


**Figure S9.** Phylogenetic structure of the initial dataset (n=490) analyzed in the study.

The tree was obtained by maximum likelihood analysis based on the final recombination-free alignment of concatenated nucleotide sequence alignments of 4,420 core genes. *Kp* ST540 Kpn0019 (accession number: SRR2098710) and a *Kp* ST101 Kp_Goe_33208 (GCF_001902435.1) were used to root the tree. Main capsular-types (KL) identified are indicated. In green the final genome dataset (n=235).
